# Promiscuous endosymbionts in deep-sea corals and crinoids are shaped by nitrogen cycling

**DOI:** 10.1101/2025.02.25.640168

**Authors:** Flúvio Modolon, Alessandro do N. Garritano, Lillian J. Hill, Gustavo Duarte, Amanda Bendia, Rafael B. de Moura, Vivian Pellizari, Torsten Thomas, Raquel Peixoto

## Abstract

Crinoids are commonly associated with corals, but whether they physiologically interact is unclear. Like corals, crinoids host symbiotic microorganisms but little is known about the crinoid microbiome. Here, we reveal the microbiomes of the deep-sea corals *Desmophyllum pertusum* and *Solenosmilia variabilis* and, notably, their associated feather star crinoid, *Koehlermetra* (of the family Thalassometridae) in the Campos Basin, Brazil. We showed that the same endosymbiotic members of the families *Endozoicomonadaceae* and *Nitrosopumilaceae* interchangeably inhabit the internal and external structures of the corals and crinoid, indicating promiscuity in symbioses. The metagenome-assembled genome of the novel and dominant *Endozoicomonas promiscua* sp. nov. suggest that these symbionts may drive dissimilatory nitrate reduction to ammonia, which could be a source of energy for ammonia-oxidizing archaea (AOA) of the family *Nitrosopumilaceae*. Thus, nitrogen cycling may determine the microorganisms that are hosted by deep-sea corals, which may be provided by their associated crinoids. These findings provide important insight about the complex ecological interactions that could lead to promiscuous symbiosis.

**Teaser:** Deep-sea corals and crinoids share symbiotic microbes that cycle nitrogen, potentially supporting each other in the nutrient-poor deep sea.

## Introduction

Symbiotic relationships between different organisms have driven the evolution and establishment of life on our planet(*1*). Multifaceted interactions, including physiological adaptations, are crucial for symbionts to thrive. Most animals, for example, rely on symbiotic relationships with microorganisms to support their physiology, ecology, and evolution(*2*). These mutualistic microbe-animal symbioses have been vital in shaping life(*3*), often providing nutritional benefits for both partners(*2*, *4*, *5*).

In general, host-microbe symbioses can be classified into two categories: first, strict symbioses, where microbial symbionts establish a relationship with a specific host and are limited to distinct niches(*6*); and, second, promiscuous symbioses, where symbionts or hosts are associated with different partners(*7–9*). Promiscuous hosts have been described in marine ecosystems. For example, the deep-sea coral *Desmophyllum pertusum* (syn. *Lophelia pertusa*) hosts highly variable microbial symbiont communities, which change with geographical location, depth, and environmental conditions(*10*, *11*). By contrast, promiscuous microbial symbionts are poorly studied, even though there are numerous examples of microbiomes being shared between different hosts (*10*, *12–16*). A few examples of well-studied tripartite host-microbe-host partnerships with promiscuous microorganism include the association of specific *Rhizobium* lineages with the roots of different legumes(*7*) and the occurrence of a specific bacterial symbiont (Candidatus *Thiosymbion*) in stilbonematine nematodes and phallodriline annelids(*8*). Such promiscuous symbionts can assume similar or identical functions in different hosts, since different hosts likely provide similar niches that are suited to a symbiont’s metabolic abilities(*17–19*). How promiscuous symbiotic relationships evolve and develop to metabolically benefit various hosts and microbes is poorly understood.

Nitrogen is a crucial nutrient for life, and its availability often limits primary production in the marine environment. Microbial communities play a vital role in driving nitrogen cycling, transforming nitrogen between various forms and making it accessible to other organisms. Key processes in this cycle include: (1) dissimilatory nitrate reduction to ammonium (DNRA), where microbes convert nitrate to ammonia(*20*, *21*), and (2) nitrification, a two-step process where ammonia-oxidizing archaea (AOA) and bacteria (AOB) first oxidize ammonia to nitrite, which is then further oxidized to nitrate by nitrite-oxidizing bacteria (NOB)^59, 60^. These microbial transformations are particularly important in the deep sea, where chemoautotrophic microorganisms become increasingly prevalent with depth(*22*). Many marine invertebrates form symbiotic relationships with nitrogen-cycling microbes, often harboring them within their tissues(*23–26*). These symbioses can provide the host with access to essential nitrogenous compounds, while the microbes benefit from a stable environment and access to host-derived nutrients. For example, microbial DNRA can be a major source of ammonia for tropical corals under dark conditions, contributing significantly to their nitrogen uptake(*25*). Understanding these intricate host-microbe relationships is crucial for comprehending nitrogen cycling dynamics and the overall functioning of marine ecosystems, especially in the deep sea where nitrogen availability can be particularly limiting.

During an oceanographic campaign in the Campos Basin (Brazil) during the winter of 2021, we observed that feather stars (crinoids, Crinoidea class, phylum Echinodermata) were often found attached to the surfaces of different species of Scleractinian corals. Crinoids are filter feeders, consuming plankton and organic particles from the water column(*27*, *28*). As both corals and crinoids host symbiotic microbes, such host interactions could facilitate a tripartite coral-microbe-crinoid partnership. Studying this intriguing and potentially promiscuous symbiotic relationship poses several challenges, including the lack of prior knowledge about the crinoid microbiome and the lack of established methods and protocols tailored to studying such potentially intricate symbiotic interactions in the deep-sea environment. We therefore explored this potential tripartite symbiotic relationship in the deep sea by investigating the potential interactions between the microbiomes of the deep sea corals *D. pertusum* and *Solenosmilia variabilis* with the feather star crinoid, *Koehlermetra* sp. that was consistently found to be associated with these two corals in the Atlantic Ocean (Campos Basin, Brazil). Our findings compare the microbiomes of crinoid and deep-sea corals and identify a new species of endosymbiont, which we call *Endozoicomonas promiscua* sp. nov. CC1, that may facilitate nitrogen metabolism in both hosts through DNRA.

## Results

### The association of a crinoid with deep sea corals

During the deep-sea expedition to the Campos Basin, we frequently found corals that were occupied by crinoids (common name: feather star) (Fig. 1a,b). The corals were identified as *Desmophyllum pertusum* and *Solenosmilia variabilis* according to their morphological characteristics and sequencing of the gene encoding cytochrome C oxidase subunit I (COI) (Supplementary Table S1). Crinoids were classified into two phenotypes, “large” and “small”, and were subdivided during sampling into body parts: large arms (or simply “arms”) plus large calyx-cirri [or simply “calyx”, which includes the large calyx (body) plus the cirri (legs) structure), and small arms (or “S_Arms”) plus small calyx-cirri (or “S_calyx”) (Fig. 1e). Crinoid specimens were identified by the diagnostic characteristics mentioned in keys for the family Thalassometridae and the genus *Koehlermetra*(*29–33*). Using sequencing of the *COI* gene, we found that both large and small phenotypes belong to the same species (100% identity), indicating that they correspond to different life stages (juveniles and adults). In addition, the crinoid *COI* sequences showed identities of 97.12 to 97.31% with *Koehlermetra porrecta*(*29*) (Supplementary Table 1). This is, to the best of our knowledge, the first record of a member of the genus *Koehlermetra* in the South Atlantic.

**Fig. 1.**
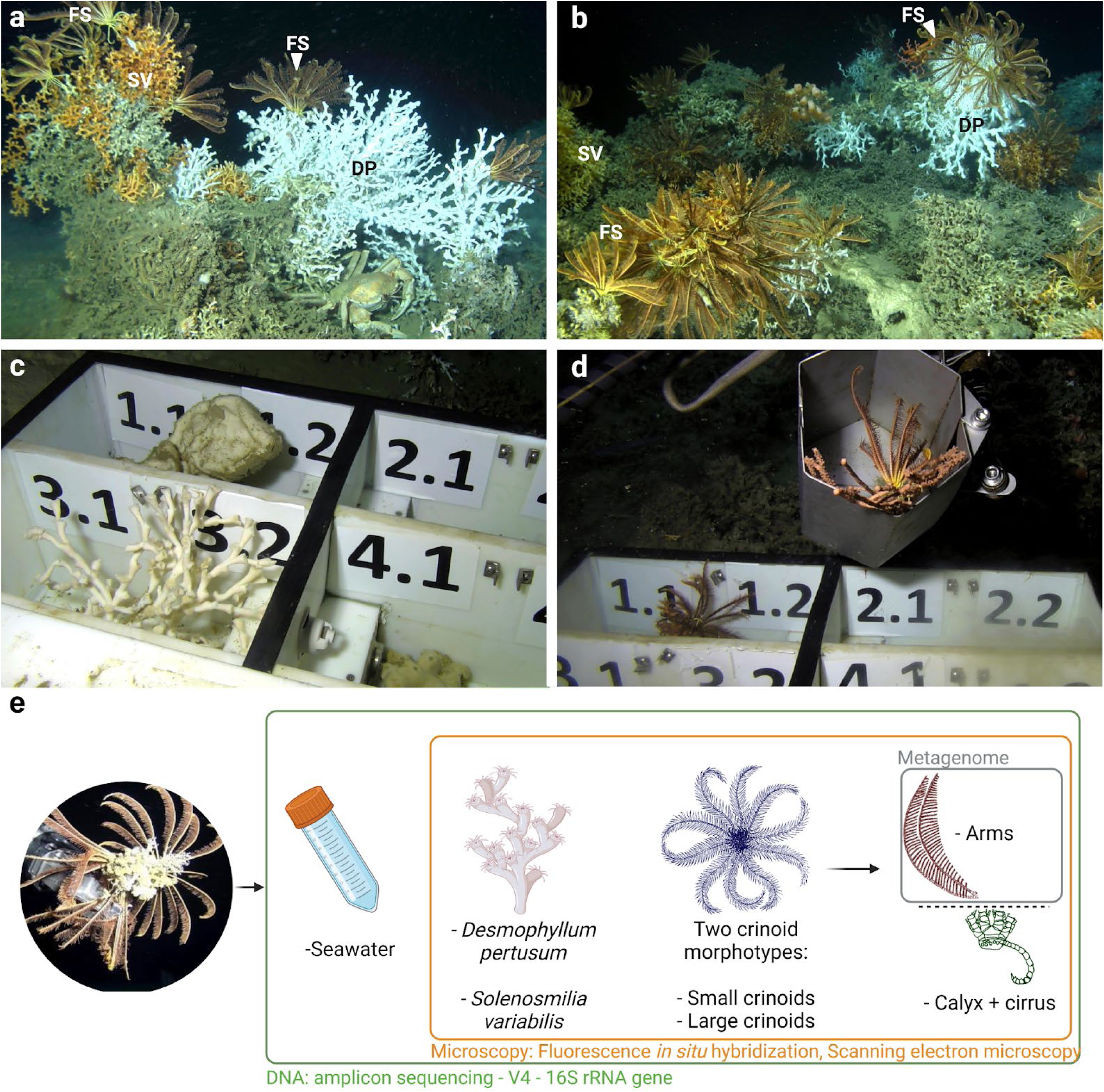
*In situ* observations and sampling of crinoid-coral associations in the Campos Basin. *In situ* photographs (a, b) showing several thalassometrid feather stars (FS) (*Koehlermetra* sp.) attached to the surface of two coral species, *Desmophyllum pertusum* (DP) and *Solenosmilia variabilis* (SV). Individual sampling boxes shown in c and d were used to collect different samples, avoiding cross-contamination. Sampling strategy and workflow are presented in e. The images were captured at a depth of 671 m in the Campos Basin, Brazil.

### Comparison of microbiomes in deep sea corals and associated crinoids

We collected coral and crinoid samples using individual sampling boxes (Fig. 1) to avoid accidental cross-contamination, and aseptically sampled their microbiomes, which were then analyzed by 16S rRNA gene amplicon sequencing. Both coral species and arms from large (adult) crinoids showed a dominance of one or a few taxa, quantified by low Pielou’s evenness (PE) values (*Desmophyllum pertusum*, PE = 0.179 ± 0.08; *S. variabilis*, PE = 0.145 ± 0.126; crinoid arms, PE = 0.387 ± 0.067), with no significant differences between corals (Fig. 2a, ANOVA, p-value = 0.9952). In a principal coordinate analysis (PCoA), the microbiomes of the large arms did not cluster with those of the other crinoid samples (large calyx, small calyx, and small arms), with statistical support for differences in terms of community composition (Sørensen dissimilarity, large arms vs. large calyx, p-value [PERMANOVA] = 0.006, p-adjusted [PERMDISP] = 0.988; large arms vs. small calyx, p-value [PERMANOVA] = 0.049, p-adjusted [PERMDISP] = 0.999; large arms vs. small arms, p-value [PERMANOVA] = 0.015, p-adjusted [PERMDISP] = 0.995).

**Fig. 2.**
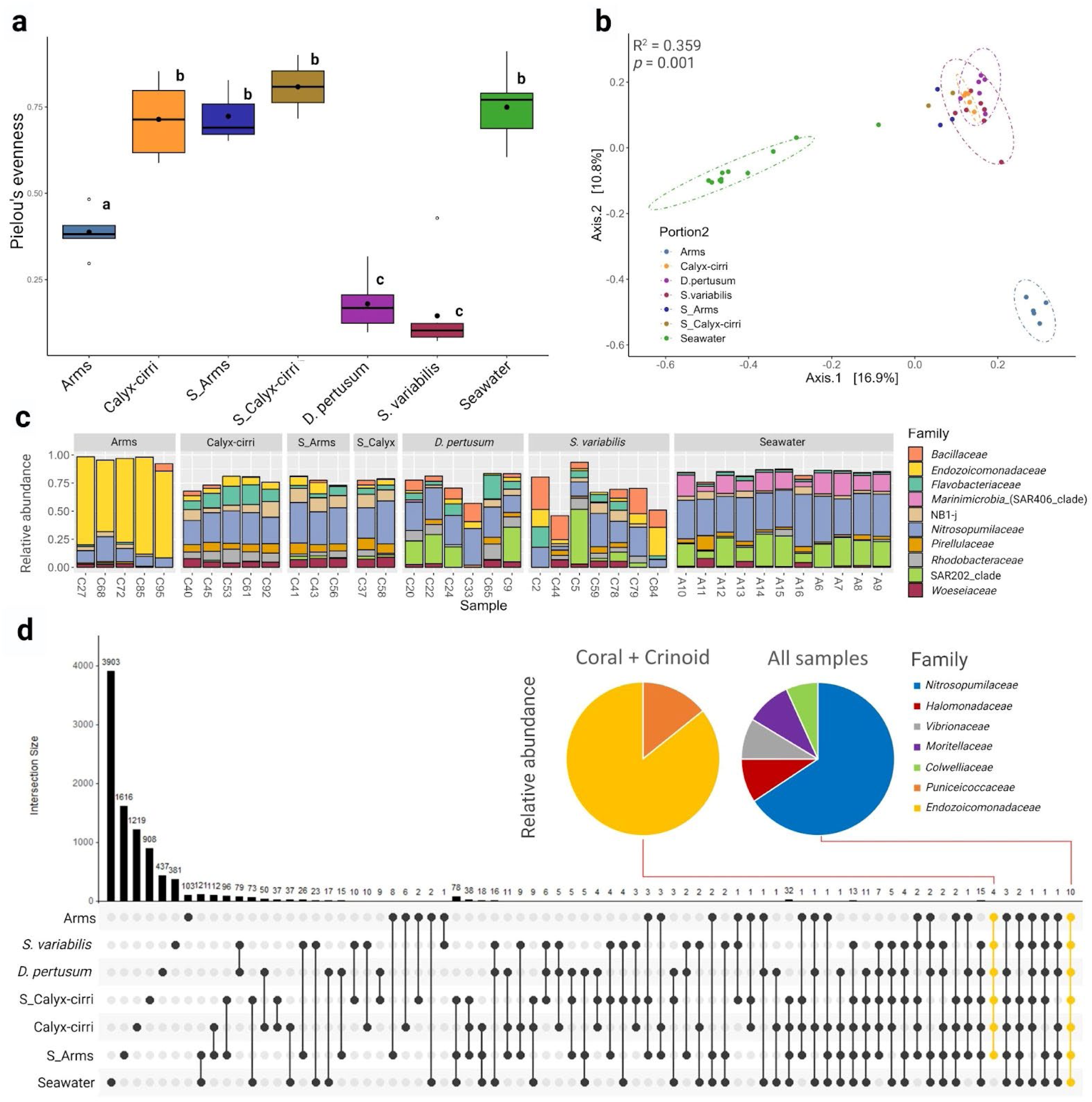
Comparison of microbial communities of crinoid and coral microbiomes. Pielou’s evenness index (a), principal coordinate analysis (PCoA) with Sørensen dissimilarity (b) for crinoids, corals, and seawater samples. Taxaplot (c) and Upset chart (d). Different lowercase letters in (a) indicate significant differences between sample groups (*p* < 0.05, ANOVA). Yellow dots in (d) highlight the intersections (shared ASVs) between all host samples (including both coral species and all body parts from large and small crinoids) and all samples (hosts and seawater), respectively. Pie charts in (d) show the relative abundances of shared ASVs between hosts (left pie chart) and all samples (right pie chart). The *x*-axis shows the ASV counts for each intersection, and the *y*-axis indicates the intersection size.

In contrast, samples of small crinoids (including small arms and small calyx) were grouped close to the samples of large calyxes, with statistical support for differences in their composition (Fig. 2b, large calyx vs. small calyx, p-value [PERMANOVA] = 0.046, p-adjusted [PERMDISP] = 0.946; large calyx vs. small arms, p-value [PERMANOVA] = 0.016, p-adjusted [PERMDISP] = 1.000). In addition, there were no significant differences between small calyx and small arms (p-value = 1.000, PERMANOVA), indicating a homogeneous microbial composition of the different body parts in the juvenile crinoids. For the coral species, no significant differences were observed in the microbial community composition of *D. pertusum* in comparison to *S. variabilis* (Fig. 2b), (Sørensen dissimilarity, p-value = 0.177, PERMANOVA). The microbial community compositions of both coral species differed from all crinoid samples (p-value [PERMANOVA] < 0.05, p-adjusted [PERMDISP] > 0.05). In addition, there were no significant differences between corals associated with crinoids and those not associated with them (Sørensen dissimilarity for *D. pertusum*, p-value = 0.400; for *S. solenosmilia*, p-value = 0.812; PERMANOVA - Supplementary Figure S1).

Amplicon sequence variants (ASVs) assigned to the family *Endozoicomonadaceae* were found associated with all host samples but were absent from seawater samples (Fig. 2c). Arms from large crinoids were especially dominated by a few ASVs (Supplementary Table S2) assigned to the family *Endozoicomonadaceae* (Fig. 2c). ASV from the archaeal family *Nitrosopumilaceae* had high relative abundances in seawater and host samples, especially in the large-crinoid calyx-cirri group. The SAR202_clade was also relatively abundant in seawater and in the majority of *D. pertusum* samples. Other microbial groups were widespread across host samples, such as NB1-j and the families *Bacillaceae* and *Woseiceaea* (Fig. 2c). Four ASVs occurred in all hosts (Fig. 2d), including three that were assigned to *Endozoicomonadaceae* and one to *Puniceicoccacea*. Together, these three *Endozoicomonadaceae* ASVs accounted for 11.84% of the total reads from 9718 observed ASVs across all samples. Ten ASVs were shared among all samples, including six assigned to *Nitrosopumilaceae*, one to the bacterial families *Colwelliaceae, Halomonadaceae*, *Moritellaceae*, and *Vibrionaceae* (Fig. 2d).

### *Endozoicomonadacea* and *Nitrosopumilaceae* ASVs are characteristic of corals and crinoid microbial communities

An indicator species analysis(*34*) showed that several ASVs are particularly prevalent and abundant in specific crinoid and coral groups. Some ASVs assigned to the family *Nitrosopumilaceae* were indicators of the groups “crinoid + *D. pertusum*”, “crinoid + seawater”, and “*D. pertusum* + *S. variabilis*” (Fig. 3a). SAR202, *Nitrospiraceae*, and *Flavobacteriaceae* ASVs, among others, were common indicators for coral species, although Stat values ranged around 0.70 for these ASVs, with specificity values ≥ 0.86 but fidelity values ≤ 0.46. ASV7675, assigned to *Spongiibacteraceae*, showed high specificity and fidelity for corals (Stat value 0.989) (Fig. 3a).. Some *Endozoicomonadaceae* ASVs were good indicators (Stat value > 0.80) of crinoids, and one ASV (8209) showed a Stat value of 0.866 for the “crinoid + *D. pertusum* + *S. variabilis*” grouping, with fidelity of 1.00 and specificity of 0.75. The mean relative abundance of ASV8209 was higher in the arms from adult crinoid samples in comparison to corals and the other crinoid samples (p-value < 0.05, Fig. 3b), with higher variance for corals. ASV8209 was detected in some, but not all, of the *D. pertusum* and *S. variabilis* replicates (Fig. 3b). These findings indicate that *Endozoicomonas* ASV8209 occurs ubiquitously in adult and juvenile crinoids, but is facultative in deep-sea corals. The ASV7675, assigned to *Spongiibacteraceae*, did not differ in mean relative abundance for *D. pertusum* and *S. variabilis* (Fig. 3c, p-value = 0.8695, ANOVA)

**Fig. 3.**
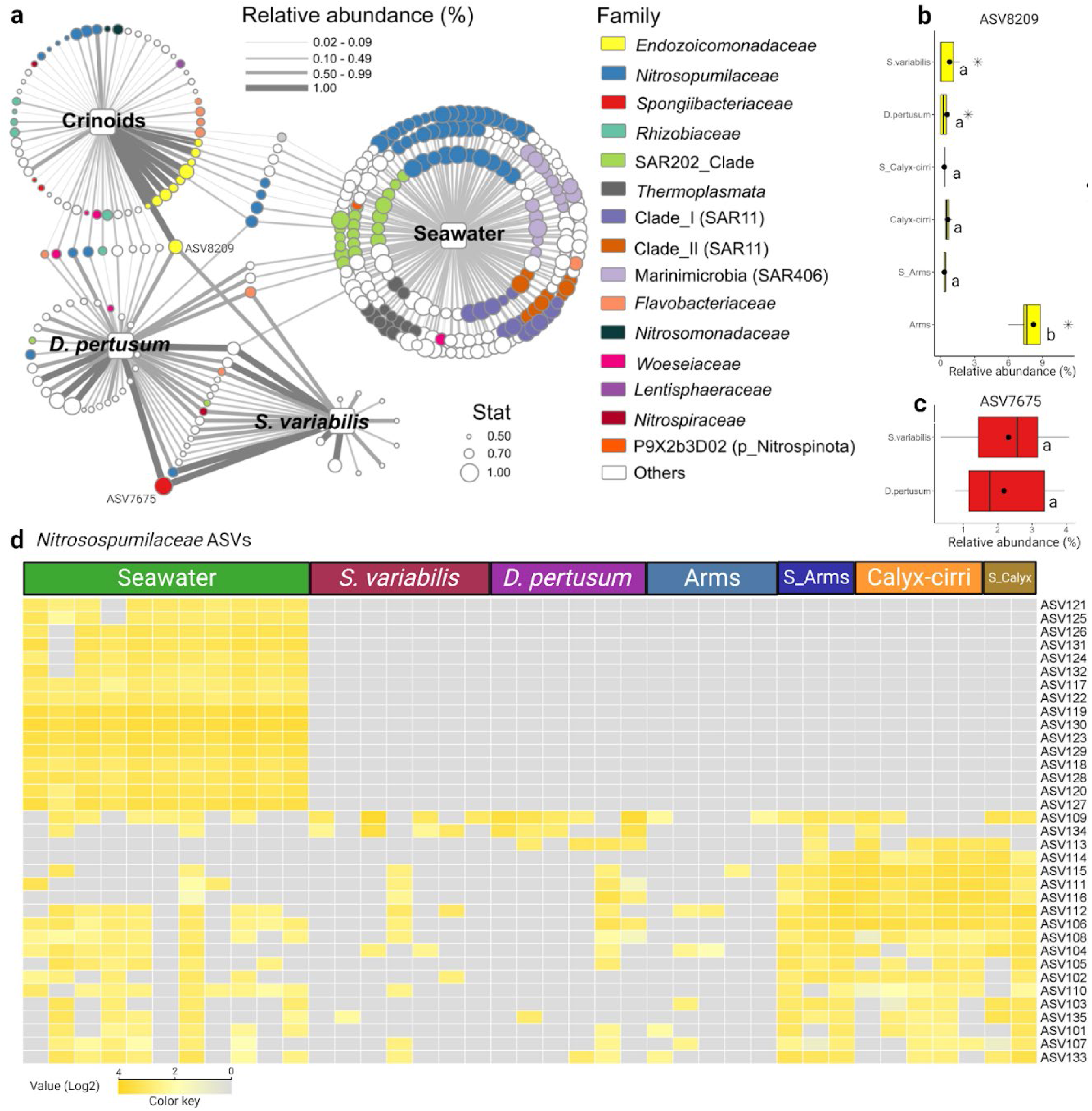
Analysis of shared and unique microbial taxa associated with crinoids and corals. a, The ASVs (circles) that characterize each sample group (represented by squares, crinoid, *Desmophyllum pertusum*, *Solenosmilia variabilis*, seawater, or the intersections between them), where colors represent the identifications at the family level, and size represents the Stat value. Stat value is the indicator value index used to predict which ASVs are good indicators for sample types. Edge thickness represents the relative abundance of the given ASV in the linked sample group. b, c, Relative abundances of ASVs 8209 (*Endozoicomonadaceae*) and 7675 (*Spongiibacteraceae*), which showed IndVal.g = 0.866 for the grouping “crinoid + *D. pertusum* + *Solenosmilia*” and 0.989 for the grouping “*D. pertusum* + *S. variabilis”*, respectively. The filled circles represent the mean value, and the asterisks (*) are outliers. d, Heatmap of Log_2_ transformed relative abundances for *Nitrosopumilaceae* ASVs. The color intensity (ranging from gray to yellow) represents the Log_2_ transformed relative abundance. Different lowercase letters in b and c indicate significant differences (p-value < 0.05) between samples.

Having found that *Nitrosopumilaceae* ASVs are indicators for “crinoid + seawater”, “crinoid + *D. pertusum*” and “*D. pertusum* + *S. variabilis*”, we selected the most abundant ASVs assigned to this taxon and evaluated their distributions across samples (Fig. 3d). Several *Nitrosopumilaceae* ASVs were observed only in the seawater samples, whereas others were found associated with hosts as well as in seawater. ASV109 appeared in both coral species and was widespread mainly in juvenile crinoid samples. ASV113 was observed in *D. pertusum* and adult crinoid calyx (Fig. 3d). This suggests that *Nitrosopumilaceae* ASV109 and ASV113 are capable of colonizing both corals and crinoids, indicating a lack of strict host specificity. The observed distribution of ASV8209 (*Endozoicomonadaceae*), ASV109, and ASV113 (*Nitrosopumilaceae*) across different host species suggests that these microbes may be promiscuous symbionts. Their ability to be associated to multiple hosts highlights the potential for complex interactions and potential cross-host microbial transfer within this deep-sea community.

### *Endozoicomonadaceae* and *Nitrosopumilaceae* are potential promiscuous endosymbionts of deep-sea corals and crinoids

To confirm the presence and analyze the spatial distribution of *Endozoicomonadaceae* and *Nitrosopumilaceae* lineages as the main symbionts shared by corals and crinoids, we utilized fluorescence *in situ* hybridization (FISH) to visualize *Endozoicomonadaceae* and ammonia-oxidizing archaea (AOA, an indicator of *Nitrosopumilaceae* species), using previously described probes(*35*, *36*). The presence of *Endozoicomonadaceae* within both crinoid and coral tissues suggests that these microorganisms are endosymbionts (Fig. 4a,b,c,d). Probes for AOA suggested that these microorganisms were widespread throughout the calyx (Fig. 4c) and scarcer in the other samples (Fig. 4a,b,d). Fluorescent signals indicating scattered microbial cells were observed in all samples. Bacilliform and cocciform microorganisms (Fig. 4e,f,g) were observed on the surface of crinoids, using scanning electron microscopy (SEM). Microbial assemblages were clearly observed inside pores, indicating that they are a potential hotspot for microbial growth and/or exchange. Across the coral surfaces, an intricate matrix of polymeric substances was observed, with cocci-shaped microorganisms predominating (Fig. 4, g). These findings reinforce the presence of *Endozoicomonas* and AOA as endosymbionts within both crinoids and corals, suggesting a potential for metabolic interactions between these microbial groups and their hosts, while also highlighting the diverse microbial communities associated with these organisms’ structures.

**Figure 4.**
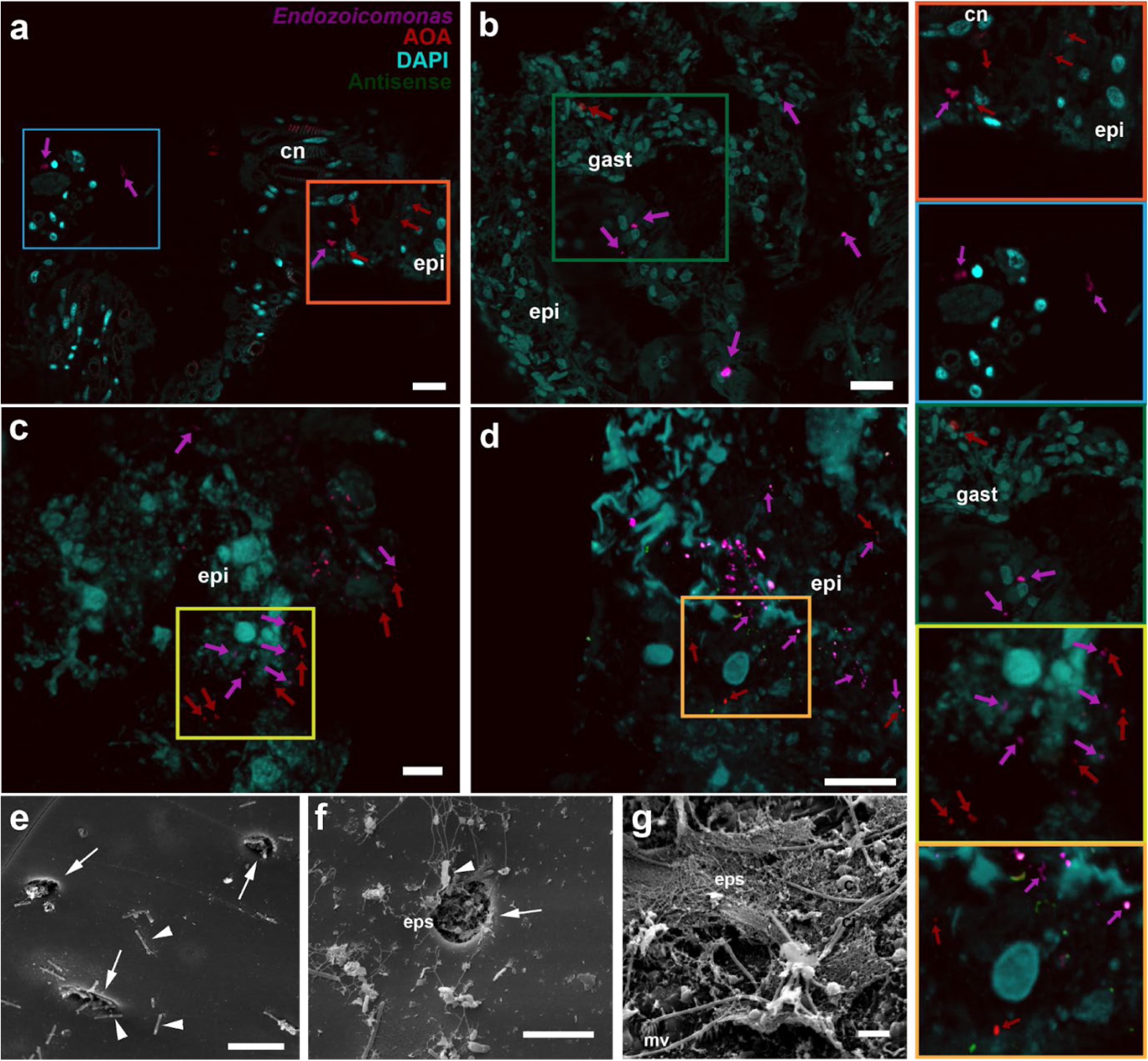
Localization of microbial cells in association with coral polyps and crinoid arms and calyx-cirri. a-d - Fluorescence in-situ hybridization (FISH) images of samples from polyps of the corals *Desmophyllum pertusum* (a) and *Solenosmilia variabilis* (b) and from the calyx-cirri region (c) and arms (d) of crinoids. Insert rectangules represent areas with higher magnification on the right of the panel, for better visualization. Blue and orange inserts are from D. pertusum, dark green insert from S. variabilis, yellow insert from the calyx-cirri region and light-orange insert from the arms of crinoids. Cyan-blue represents DAPI, red represents ammonia-oxidizer-archaea specific probe conjugated with Alexafluor-647 (AOA, red arrows), purple represents *Endozoicomonas* spp. probe conjugated with Alexafluor-594 (purple arrows), and green represents Non-Eubac and Non-Archaea probes (conjugated with Alexafluor-488) to account for unspecific binding. Cn = Cnidocytes; Epi = Epidermis; Gast = gastrodermis. See methods for specific information about probe designs and FISH protocols. Bars = 10 µm. Other sections including alternative fluorophores and antisense probing (Non-Eubac and Non-Archaea) images can be found in the Supplementary Figures S3, S4 and S5. e-g - Scanning electron micrographs of the surface of coral-associated crinoid calyx and coral polyp. e,f - Pores that connect the epidermis with the crinoid skeleton (arrows), bacilli (b), and coccus (arrowhead) cells, as well as EPS (especially in the pore spaces). g - coral cell structures – microvilli (mv) and cilia (c) – as well as a high abundance of EPS and coccus cells. Bars = 5 µm.

### *Endozoicomonas* spp. metabolize nitrogen

The high abundance and promiscuous associations of *Endozoicomonas* species with deep-sea corals and crinoids indicates these microbes have a potentially important role. However, there is limited knowledge about their functional traits. To investigate the metabolism and potential symbiotic role of *Endozoicomonadaceae*, we conducted shotgun metagenomics followed by binning of the adult crinoid arm samples, which were dominated by this taxon (particularly ASV8209). A metagenome-assembled genome (MAG) from the genus *Endozoicomonas* sp. (MAG-CC1) was binned with a predicted completeness of 96.17%, a bin size of 5.37 Mb, 2.87% predicted contamination, and a GC content of 47%. Genome annotation revealed that this *Endozoicomonas* MAG has a complete pathway for dissimilatory nitrate reduction to ammonia (DNRA), in addition to other genes related to nitrogen metabolism, including the nitrate/nitrite transporter NrtP (See Supplementary Table S3). Phylogenomic analysis showed that MAG-CC1 was grouped closer to *E. elysicola* DSM22380 than to other genomes (Fig. 5). The difference in GC content between MAG-CC1 and *E. elysicola* DSM22380 was 0.49% and the *in silico* DNA-DNA hybridization value was 25.10%. In addition, the mean nucleotide identity (ANI) and the mean amino acid identity (AAI) between MAG-CC1 and *E. elysicola* DSM22380 were 80.50% and 85.37%, respectively. According to the proposed threshold values for taxonomic classification of uncultured prokaryotes(*37*) using ANI/AAI values (45–65%, 65–95%, 95–100% for family, genus, and species, respectively), the genomes of MAG-CC1 and *E. elysicola* DSM22380 belong to different species of the same genus. Furthermore, we propose a new species epithet for MAG-CC1, *Endozoicomonas promiscua* sp. nov. CC1, according to SeqCode rules and recommendations(*38*).

**Figure 5.**
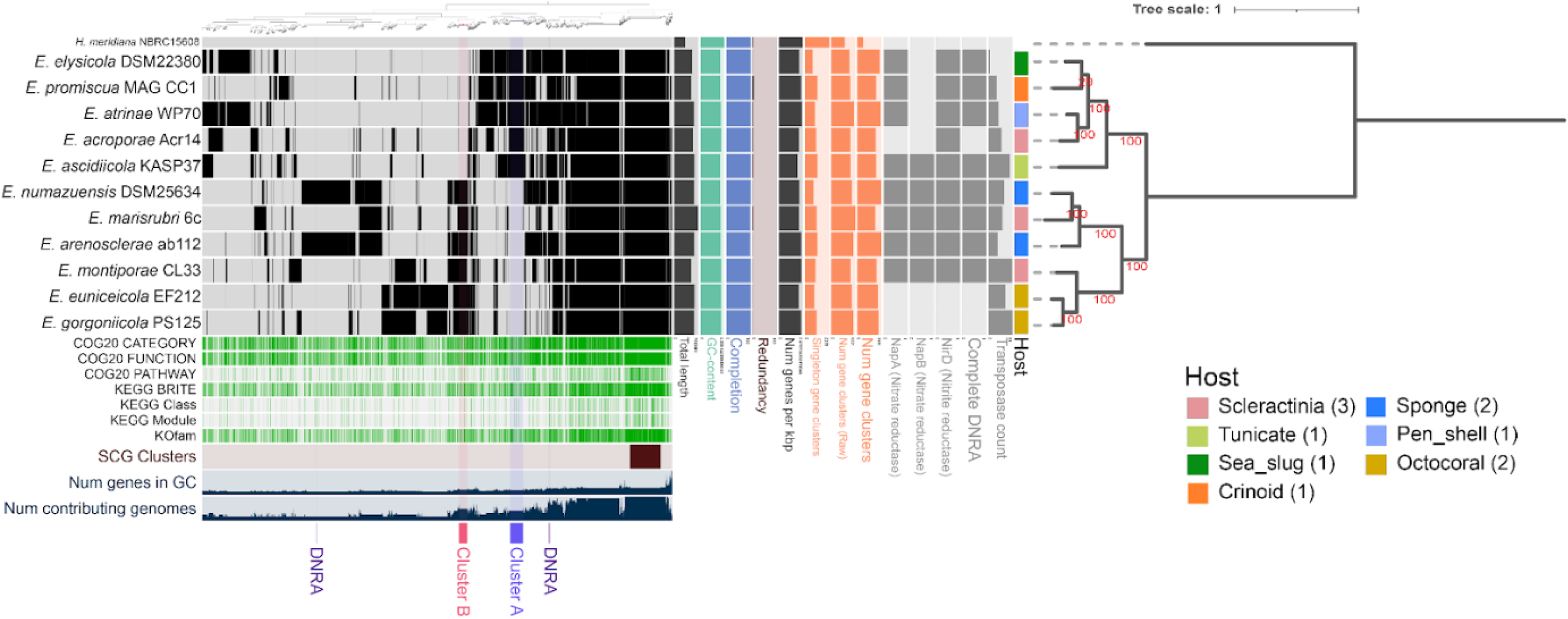
Pangenome and phylogenomic analysis for *Endozoicomonas* genus. For the pangenome analysis, we compared 10 high-quality *Endozoicomona*s genomes (downloaded from NCBI and RAST) plus MAG-CC1 from this study. Overlapping coloured bars indicate gene clusters shared between genomes. The number of transposase counts refers to dereplicated coding sequences observed on each genome. Nitrate reductase (genes napA and/or napB) and Nitrite reductase (gene nirD) compose the Dissimilatory Nitrate Reduction to Ammonium (DNRA) pathway. The figure was generated in Anvi’o and the genomic annotations and search for DNRA genes were carried out using Rapid prokaryotic genome annotation (Prokka 1.14.6), the database of Clusters of Orthologous Groups (COG) and KEGG. The scale bar corresponds to 0.05 substitutions per nucleotide position. The numbers on nodes refers to the levels of bootstrap support (%). Clusters A and B represent the genes distinguishing the two monophyletic groups formed in the tree.

Pangenome analysis comparing 10 high-quality *Endozoicomonas* spp. genomes and *E. promiscua* CC1 revealed that these genomes are highly variable in gene content and size, with a small set of core genes (Fig. 5), in agreement with previous findings by Neave and collaborators(*39*). Despite this high variability, genomic annotation revealed that eight of these genomes harbor the complete pathway for DNRA, and nine have at least one component of DNRA (e.g., genes for nitrate reductase or nitrite reductase). In addition, a variable count of transposases ranging from 4 to 65 was observed in the genomes (Fig. 5).

The phylogenomic tree revealed two monophyletic groups for *Endozoicomonas* that were distinguished according to core gene sets in: MAG-CC1, *E. elylisicola* DSM22380, *E. atrinae* WP70, *E. ascidiicola* KASP37 and *E. acroporae* Acr14 (Cluster A); and *E. numazuensis* DSM25634, *E. marisrubri* 6c, *E. arenoscleare* ab112, *E. montiporae* CL33, *E. euniceicola* EF212 and *E. gorgoniicola* PS125 (Cluster B) (Fig. 5). Genes for different amino acid biosynthesis differentiate Cluster A and B. Exclusive to Cluster A were genes for glycine, serine, threonine, valine, leucine, isoleucine, and lysine biosynthesis (Supplementary Table S4). In contrast, Cluster B exclusively contained genes for arginine, proline, and histidine biosynthesis (Supplementary Table S4). Although both clusters contained genes for cysteine and methionine biosynthesis, the specific genes differed between clusters. Overall, these findings reveal that *Endozoicomonas* genomes are highly diverse, particularly in terms of nitrogen metabolism genes, and that the novel species identified here, *E. promiscua* sp. nov. CC1, possesses the complete pathway for DNRA, suggesting their potential role in nitrogen cycling within both crinoid and coral holobionts.

Despite extensive efforts, obtaining *Nitrosopumilaceae* MAGs was challenging. However, the extensive and well-known metabolic traits of *Nitrosopumilaceae* available in the literature, mainly related to the nitrogen cycle, provided a well-grounded discussion of such interactions.

### Nitrogen-metabolising microbes in deep-sea coral and crinoid microbiomes

In addition to the key promiscuous symbionts identified in this study (*Endozoicomonadaceae* and *Nitrosopumilaceae*), we also investigated other taxa associated with deep-sea corals and crinoids that are known to be involved in the nitrogen cycle (Fig. 6). For example, genomic, metagenomic, and transcriptomic evidence has shown that the members of the SAR202_clade have roles in nitrate uptake and nitrate reduction to ammonia(*40*, *41*). The genus *Nitrospira* (family *Nitrospiraceae*)(*42*) and the family *Nitrospinaceae*(*43*) are known as nitrite-oxidizing bacteria (NOB), generating nitrate. Interestingly, both these NOBs taxa were missing from adult crinoid arm samples. Members of the family *Kiloniellaceae* are known to use nitrate as the final electron acceptor, reducing it to N_2_O(*44*, *45*). The family *Woeseiaceae*, in contrast, are known to reduce nitrite to N_2_O, although their role in the complete denitrification pathway to generate N_2_ is still unknown(*46*). In addition, the family *Rhizobiaceae* is well known for nitrogen fixation in soil(*47*) and was more recently found in deep-sea environments(*48*, *49*). Fixed nitrogen could be used by ammonia-oxidizing archaea and bacteria (AOA and AOB) such as those from the *Nitrosopumilaceae* family. The presence of those taxa in both coral and crinoids reinforces that nitrogen cycling may play a crucial role for the hosts.

**Fig. 6.**
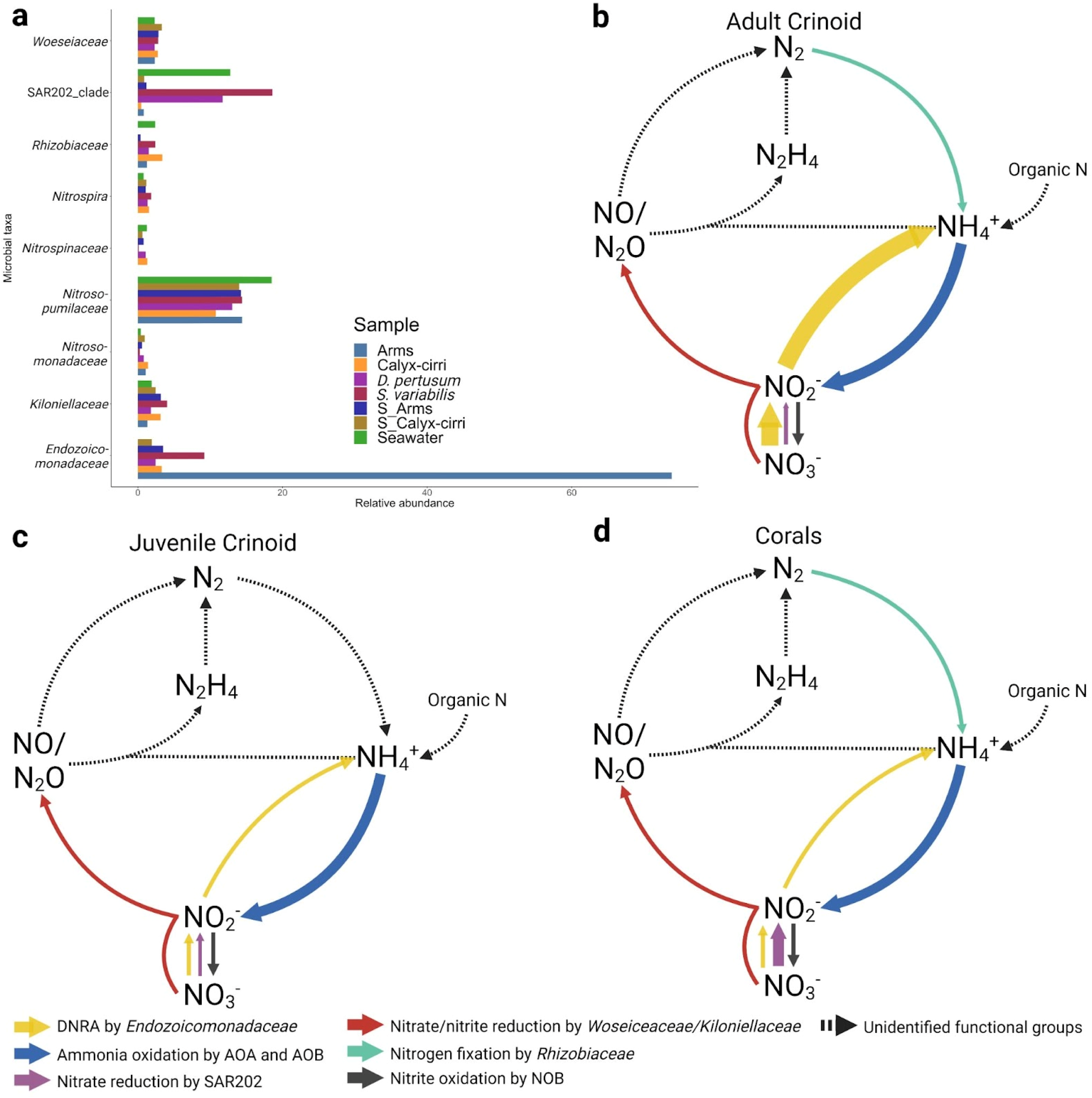
Relative abundance and predicted functional roles of nitrogen-cycling microbial taxa associated with crinoids and deep-sea corals. The histogram in a shows the relative abundances of the main taxa that have been reported as involved in nitrogen cycling in deep-sea corals and crinoids. The proposed pathways for nitrogen recycling in the hosts as well as the potential microbial taxa related to each step are represented in b for adult crinoids, c for juvenile crinoids, and d for corals. The thickness of the arrows indicates whether the groups are more or less abundant. The cycle was adapted from Jetten (2008).

We also measured the levels of nitrate, nitrite and ammonium in addition to dissolved organic carbon (DOC) in the seawater surrounding the hosts (Supplementary Table S5). Spearman correlation analysis between the 15 most abundant microbial taxa (relative abundance at family level) and these measurements showed positive correlation (Spearman coefficient = 0.3, p-value = 0.018) between *Endozoicomonadacea* and DOC. A negative correlation between nitrate and *Endozoicomonadacea* (Spearman coefficient =-0.3, p-value = 0.123) was also observed (Supplementary Fig. S2). This reinforces that *Endozoicomonadaceae* representatives contribute to nitrate reduction.

## Discussion

This study has characterized the microbiomes of two species of deep-sea coral (*D. pertusum* and *S. variabilis*) and their associated crinoid (*Koehlermetra* sp.) from the Campos Basin, Brazil. This is also the first description of a crinoid microbiome. We identified several microbial species that are common to the corals and crinoids, including a newly identified species, *Endozoicomonas promiscua* sp. nov. CC1. Additional analyses indicate that this and species of *Nitrosopumilaceae* may be promiscuous endosymbionts that metabolise nitrogen. The presence of nitrogen-metabolizing microorganisms in the microbiomes of the corals and the crinoid suggest potential microbially redundant functioning in animals from different phyla in this environment. Based on the dominance of *Endozoicomonadaceae* in the crinoid and the presence of specific ASVs found in both crinoids and corals, we hypothesise that crinoids may act as a consistent reservoir of *Endozoicomonas* spp. that can perform DNRA, which could contribute to nitrogen cycling. Through DNRA, *Endozoicomonas* spp. could provide an additional source of ammonia under N-limited conditions for the abundant *Nitrosopumilaceae*, as well as for the host in this system (Fig. 7). Overall, our results suggest that the microbiomes in deep-sea corals and their associated crinoid seem to be shaped by nitrogen cycling. The key functional aspects associated with this symbiosis and how it evolved between such different animals co-existing in the same extreme environment require further investigation.

**Fig. 7.**
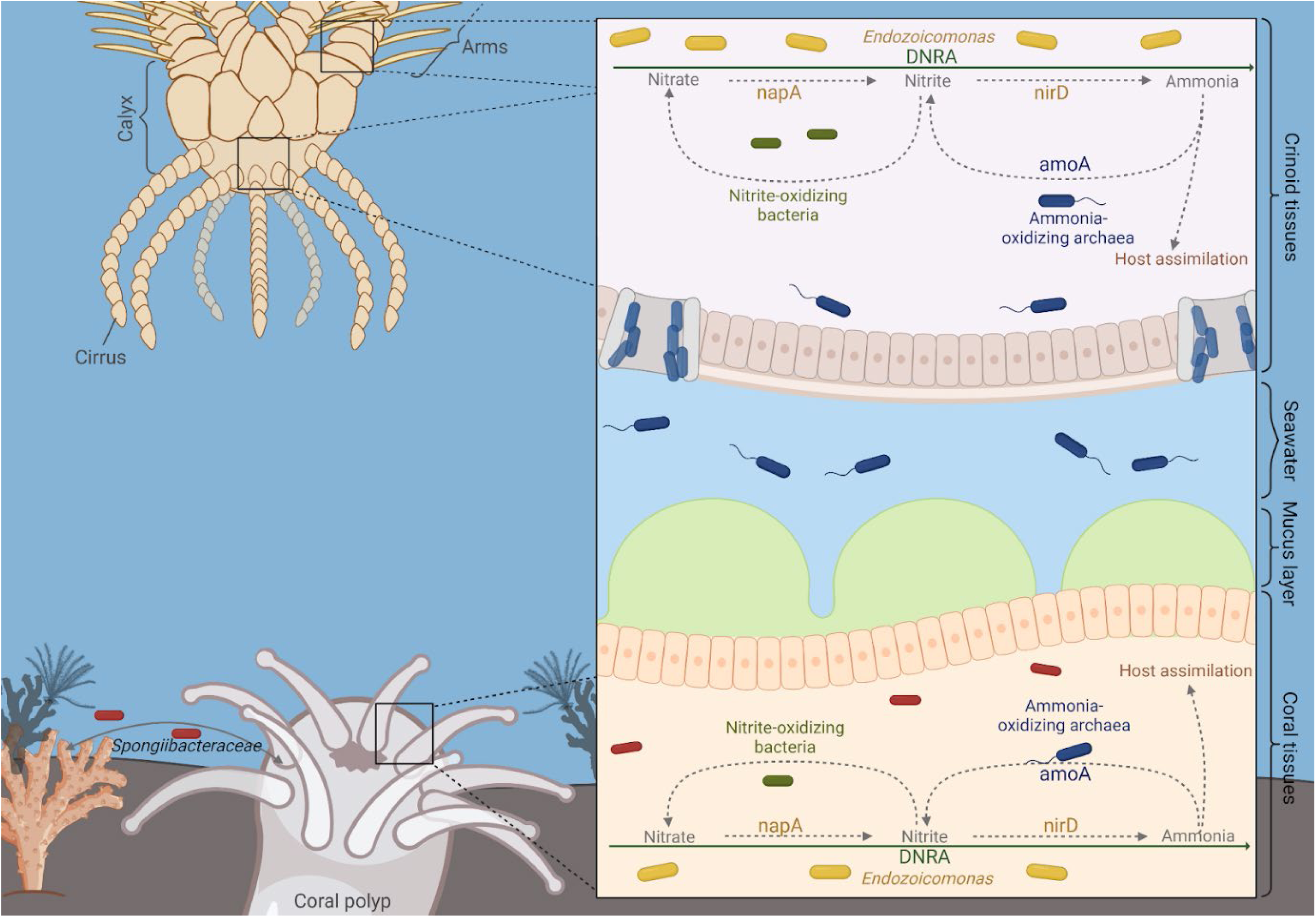
Graphical overview showing the hypothetical microbial switching between corals and the crinoid, besides the proposed metabolic interaction between AOA and *Endozoicomonadacea*. The pathway for dissimilatory nitrate reduction to ammonia (DNRA) is catalyzed by enzymes encoded by *nirD* and *napA* genes found in *Endozoicomonas* sp. MAG-CC1. The ammonia is oxidized by ammonia-oxidizing archaea, which harbor the amoA gene. Nitrite-oxidizing bacteria convert nitrite to nitrate.

The presence of AOA and other organisms involved in nitrogen cycling, in addition to the complete DNRA pathway in *Endozoicomonas*, suggest microbial contributions to nitrogen cycling in the crinoid-coral system. More specifically, we found that the most abundant families potentially involved in nitrogen cycling are the same organisms for the corals *D. pertusum* and *S. variliabilis*, as well as for both adult and juvenile crinoids. The AOAs of the family *Nitrosopumilaceae*, for example, were found associated with all host samples. In addition, the AOB assigned to the family *Nitrosomonadaceae* was also observed in all hosts, although at lower relative abundances. The consistent presence of these microorganisms in our samples, including our observations suggesting that they are endosymbionts, indicate potential ammonia oxidation within the host tissues. Other studies have also found AOA to be associated with *D. pertusum*(*50, 51*) and have shown ammonia oxidation to nitrate by isotope analysis(*23*). However, nitrification had not been previously explored in *S. variabilis* and deep-sea crinoids.

The nitrate generated could, in turn, be assimilated by the hosts(*52*). However, Middelburg et al.(*23*) demonstrated that nitrification contributes to less than 1% of the ammonia consumption of *D. pertusum*, and ammonia is mostly assimilated into organic forms by the coral host. Alternatively, nitrate could still be assimilated by coral-associated microorganisms that harbor nitrate reductases and/or the complete DNRA pathway. DNRA is a two-step pathway that plays a crucial role in nitrogen cycling by generating ammonia from NOx, which serves as a potential source for ammonia oxidation or can be incorporated into biomass. In contrast, the gaseous by-products of denitrification (NO, N_2_O, and N_2_) cannot be readily assimilated and are prone to be lost to the environment(*53*). The complete pathway for DNRA was found in *Endozoicomonas* sp. MAG-CC1 genome and in 72.7% of the previously published *Endozoicomonas* spp. genomes analysed here (Figure 4). Its activity could therefore provide an additional source of ammonia for the corals and/or their symbionts, although further studies are needed to confirm the expression of DNRA genes in *Endozoicomonas*.

Although we detected members of the family *Endozoicomonadacea* ubiquitously distributed in the crinoid, they were particularly dominant in the arms of adult crinoids (Fig. 2). As crinoids are filter feeders, the primary function of their arms is to capture food particles by water filtration(*27*, *28*), which may also facilitate the uptake of nitrate from the surrounding seawater. Alternatively, the nitrite produced by AOA could be directly used as an electron acceptor by *Endozoicomonas* sp., generating ammonia as a byproduct of NirD activity (Figure 4), in a shorter pathway(*54*).

Other taxa that can be involved in the nitrogen cycle were also found in lower relative abundances in comparison to *Endozoicomonadaceae* and AOA. These include ASVs assigned to *Rhizobiaceae* found in coral species and adult crinoids, but absent in the juveniles. They could contribute to nitrogen fixation within the hosts, as observed by Middelburg et al.(*23*) in *D. pertusum*. The SAR202 clade, which can harbor genes for nitrate and nitrous oxide reductases(*40*, *41*), was also observed in all samples, being more abundant in juvenile crinoids. ASVs assigned to the families *Woeseiaceae* and *Kiloniellaceae*, which are capable of using nitrite and nitrate, respectively, as final electron acceptors, reducing them to N_2_O, which might “gas out” from the system(*44–46*), were also found.

A meta-analysis using data from different 16S rRNA gene amplicon sequencing studies of deep-sea coral has indicated that the presence of *Endozoicomonas* spp. in stony and soft corals is variable(*10*). More recently, Kellogg and Pratte(*55*) showed that *Endozoicomonas* spp. are associated with four different deep-sea coral species (*D. pertusum*, *D. dianthus*, *Acanthogorgia aspera*, and *A. spissa*), and found an apparent dominance and diversity in samples collected below 1000 m in the Gulf of Mexico, highlighting a trend of DNRA-performing bacteria to be associated with corals in nitrogen-limited deep-sea environments. In fact, DNRA is upregulated by high environmental carbon-to-nitrogen (C/N) ratios(*56*, *57*). Our results show a positive correlation between *Endozoicomonadaceae* abundance and dissolved organic carbon concentrations, in contrast to the negative correlation with nitrate concentration, corroborating the preference for DNRA under nitrogen-limited sites. These results reinforce the potential contribution of the detected DNRA pathway for an additional input of ammonium in the system. Other evidence also indicate that DNRA is preferred in deep-sea conditions, in contrast to shallow waters(*58*). For example, genes for DNRA have been identified in symbionts associated with other deep-sea animals including tubeworms(*59*) and snails(*60*). It is unclear, however, why *Endozoicomonas* spp. would prefer anaerobic respiration with DNRA rather than aerobic in the system observed here. Recent evidence has shown that cells of *Endozoicomonas* spp. are membrane-encased inside coral tissues(*61*), which could create a specific anoxic or low-oxygen microenvironment. Alternatively, DNRA could occur under aerobic conditions, as demonstrated previously(*62*, *63*).

*D. pertusum* appears to be unique in its ability to adapt to different conditions, living under a wide range of depths, temperatures, and geographic conditions(*64–66*), thus, microbiome plasticity may be important to support this flexibility. Considering that *Endozoicomonas* spp. were ubiquitous in the crinoids and facultative for both coral species, we suggest that they have a promiscuous lifestyle by swapping hosts in this system and thus contributing to microbiome plasticity. Corals therefore may recruit *Endozoicomonas* spp. during biotic or abiotic changes in the environment that lead to changes in nitrogen turnover, and the associated crinoids could act as an inoculation source of these microorganisms. Additionally, crinoids could increase the amount of ammonia - by filtration for feeding behaviour - in the vicinity of their associated coral, which could stimulate the nitrification process in corals and provide a niche for DNRA bacteria.

The detection of shared microbes between deep-sea coral species and between crinoids and corals indicates that specific symbionts are capable of establishing relationships with different hosts. This promiscuity could be explained by the ecological-fitting premise(*19*), which means that a symbiont has metabolic capacities and niche requirements that can be provided by different hosts(*17–19*). These promiscuous symbionts are thought to have larger than average genome sizes in comparison to strict symbionts owing to the genetic isolation that results in genome reduction(*19*, *67–69*). Genomes of *Endozoicomonas* spp. vary in size and genetic content, as shown here and by Neave et al.(*39*), indicating their plasticity to adapt to different niches. MAG-CC1 has an intermediate genome size of 5.37 Mb. In addition, the enrichment of transposases within *Endozoicomonas* spp. genomes (Fig. 4) could be a mechanism to expediently acclimate to new hosts or take advantage of different niches(*39*, *70*). Furthermore, the presence of different sets of core genes related to amino acid metabolism, which differentiate two monophyletic groups of *Endozoicomonas*, suggests that these symbiotic groups may have distinct functional specificities(*12*). This implies that the different clusters possess varying amino acid biosynthesis capacities, potentially complementing the diverse profiles of host auxotrophy.

A *Nitrosopumilaceae* ASV was also shared between deep-sea corals and crinoid hosts, and some genomic adaptations to a symbiotic lifestyle could include an increased GC content(*71*); enrichment of genes related to replication, recombination, and repair; and reduction of genes related to biogenesis of the cell wall/membrane/envelope(*72*). In addition, other microorganisms may be exchanged between different hosts, such as the family *Woeseiaceae*, which was found to be associated with both *D. pertusum* and crinoids. *Spongiibacteraceae* AS7675 was also ubiquitous and promiscuous between the two coral species. Previous studies identified this family as dominant in several samples of *D. pertusum*(*15*) and widespread among gorgonians(*73*), although the function of *Spongiibacteraceae* for corals remains unclear.

As crinoids are filter-feeders, *Nitrosopumilaceae* ASVs retrieved from their tissues may reflect a combination of tissue-associated lineages (here demonstrated with FISH) and those derived from the crinoid’s feeding behaviour, once the seawater samples were enriched by such taxon. Future investigations employing techniques such as NanoSIMS and transcriptomics could provide further insights into the functional role of *Nitrosopumilacaeae* in the crinoid microbiome.

Overall, this study provides a crucial first step in understanding the complex interactions between deep-sea corals, crinoids, and their associated microbiomes. The discovery of promiscuous nitrogen-cycling symbionts, such as *Endozoicomonas promiscua* sp. nov. CC1 and specific *Nitrosopumilaceae* ASVs, raises intriguing questions about the role of these microbes in facilitating nutrient exchange and supporting the holobiont in the oligotrophic deep-sea environment. Future research should focus on investigating the functional significance of these symbioses, including the expression and activity of nitrogen metabolism genes, the potential for nutrient transfer between host and symbiont, and the impact of environmental factors on these interactions. Furthermore, exploring the broader ecological implications of microbial sharing within deep-sea communities, particularly in the context of increasing anthropogenic pressures on these fragile ecosystems, will be crucial for conservation efforts. Ultimately, deciphering the complex relationships between deep-sea organisms and their microbial symbionts is essential for understanding the resilience and functioning of these unique ecosystems in a changing ocean.

## Materials and Methods

### Deep-sea Oceanographic campaign

The study was conducted during the ProBioDeep oceanographic campaign, in August-September 2021, in the Campos Basin, Southwest Atlantic, Brazil. The DP2 research vessel Fugro Aquarius was equipped with two FCV 3000 working class remotely operated vehicles (ROVs) capable of diving up to 3000 metres. The ROVs were equipped with 5-axis manipulators and underwent custom-made modifications developed by our team, to enable sample transportation in their lower frames, known as the individual “BioDrawers”. The remotely operated BioDrawer consisted of a 20mm thick high-density polyethylene (HDPE) drawer designed to provide thermal insulation, and contained 12 dividers to optimize the simultaneous collection of samples while preventing cross-contamination.

### Sample processing, DNA extraction and sequencing

Different coral species and crinoids and their physical association were observed at depths of 670 to 830 m in the Campos Basin. Representatives of two coral species (*Desmophyllum pertusum* and *Solenosmilia variabilis*) and two crinoid morphotypes (“large crinoids” and “small crinoids”) were collected in 3–7 replicates at four oceanographic stations within the same area of the Campos Basin (Supplementary Table S5). The animals were collected randomly across the four sites aiming to increase biological variability. A single crinoid specimen per associated coral colony was collected. Similarly, a single coral fragment from the same coral colony was collected. Six *D. pertusum* samples were collected, while seven *S. variabilis* samples were sampled. The coral portions immediately associated with the crinoids were sampled. In cases where the corals were not associated with crinoids, a random portion of the colony was sampled. We obtained five (n=5) large crinoids and three (n=3) small crinoids. The crinoid body was immediately separated into the upper portion (named “arms” for large crinoids and “small arms” for small ones) and its lower portion (“calyx-cirri” for large crinoids and “small calyx-cirri” for the small ones). During the sampling processing, one replicate referred to the “small calyx-cirri” sample was lost. A total of eleven seawater samples (10 L each) were collected around the animals within this region, composed by 2-3 replicates per sampling site, and filtered onto 0.22 µm membranes (MF-Millipore, Ireland).

We acknowledge the constraints in the statistical confidence interval for the group “small calyx-cirri” where sample sizes are limited to n=2. Given the unique ecological context of our research, obtaining a larger number of replicates in certain groups was logistically challenging. Due to the singularity and pioneerism of this study, in addition to the challenges faced in the sampling processes, those samples are arguably considered rare and restricted. These limitations underscore the exploratory nature of this particular data and provide a candid framework for interpreting the “small calyx-cirri” results.

The animals and filter membranes were immediately stored in liquid nitrogen and transported to the laboratory, where they were kept at –80 °C until the DNA extractions. Total DNA was extracted from the coral, crinoid, and seawater samples, using the DNeasy PowerSoil Pro Kit (Qiagen, Hilden, Germany) according to the manufacturer’s protocol. Corals and crinoids were macerated with a mortar and pestle, and 0.5 g of the resulting slurry was used for extractions. Membranes from water sampling were fragmented using sterile scissors before the extractions.

The V4 region of the 16S rRNA gene was amplified by PCR with the primers 515F (5’-GTGYCAGCMGCCGCGGTAA-3’) and 806R (5’-GGACTACNVGGGTWTCTAAT-3’), which were appended with universal Illumina tags. The PCR thermocycling conditions involved an initial denaturation step of 3 min at 94 °C, followed by 32 cycles of 45 s at 94 °C, 1 min at 50 °C, and 90 s at 72 °C, with a final 10-min extension step at 72 °C. The cleaned PCR samples were used as input for indexing PCR reactions to add unique dual indexing tags (IDT) to each sample. An Integra Viaflo 96 was used to set up each PCR plate. All samples were normalised using a Sequelprep kit (Thermo) and eluted in 20ul of elution buffer. An equal volume was used for pooling. The pool was bead cleaned using Ampure XP beads. The cleaned pool was quantified using tapestation and qubit to calculate molarity before sequencing. The resulting amplicons labelled with barcodes were combined and sequenced with the Illumina Miseq platform, in accordance with the manufacturer’s guidelines, at the Ramaciotti Centre for Genomics (UNSW).

Sanger sequencing was carried out for the gene of the cytochrome oxidase subunit 1 (COI), using the primers LCO1490 5′–GGTCAACAAATCATAAAGATATTGG–3′ and HCO2198 5’-TAAACTTCAGGGTGACCAAAAAATCA-3’ for corals and the primer pair FsCOI 5′-AGTCGTTGGTTGTTTTCTAC-3′ and COI 3′R 5′-CAATGAGTAAAACCAGAA-3′ for comatulid crinoids (Helgen and Rouse, 2006).

The final data set consisted of 39 samples, including 13 corals (6 *D. pertusum*, 7 *S. variabilis*), plus large crinoids and small crinoids, which were subdivided into 5 large arms, 5 large calyx-cirri, 2 small arms, and 3 small calyx-cirri, and 11 seawater samples.

### Metagenomic: extraction, sequencing, assembly, and annotation

To enrich microbial cells from host tissue for DNA extraction, we employed a series of filtration and centrifugation steps. Initially, approximately 0.8 g of crinoid arms was rinsed with sterile calcium-and magnesium-free artificial seawater (CMFASW), after which the sample was homogenized with 2 ml of sterile CMFASW in a mixer. The resulting homogenate was then filtered through a 100-μm filter, and the filtrate was centrifuged twice at 300 *g* and 4 °C for 15 min each to decant debris. The supernatant was subsequently filtered twice through a 20-μm filter (Nylon Net, Merck Millipore, Ireland) and then through a 3-μm membrane (MCE, MF-Millipore, Ireland). Finally, the resulting filtrate was centrifuged at 15,000 *g* and 4 °C for 15 min to pellet the microbial cells, which were then resuspended in EDTA and snap-frozen for eventual DNA extraction. DNA extractions were performed using DNeasy PowerSoil Pro Kit (Qiagen) according to the manufacturer’s protocol, except for the fast-prep step, for which we used speed 4 for 30 s.

Library preparation was performed using the Illumina Stranded mRNA Prep Ligation kit with modification of starting at the fragmentation step with 1ng input and 15 PCR cycles. Illumina IDT for Illumina RNA UD Indexes were used to uniquely identify samples. Libraries were quality-checked using the ThermoFisher Qubit 4.0 fluorometer (dsDNA HS assay) and the Perkin Elmer GX Touch HT (High Sensitivity DNA assay) and sequenced on a Illumina NovaSeq X Plus platform (2 x 150bp). All the crinoid-associated reads had the adapters removed using Trimmomatic (v0.38)(*74*). The resulting reads were assembled using metaSPAdes (v3.12.0)(*75*) and mapped in an all-versus-all manner using Bowtie (v2.3.5.1)(*76*) to obtain an indexed BAM file for differential coverage estimation. Samtools (v1.10)(*77*) was used to generate an indexed SAM file for estimating coverage information. Binning was performed using MetaBAT (v2.12.1)(*78*), MyCC (v20170301)(*79*), and SemiBin (v1.3.0)(*80*), and refined with Metawrap (v1.3)(*81*). A first screening for taxonomy was assigned to the obtained MAG based on the Genome Taxonomy Database (GTDB, release 207)(*82*) using GTDB-Tk (v2.1.1)(*83*), which classified the MAG based on placement in a reference tree inferred using a set of 120 bacterial gene markers, using a combination of FASTANI and pplacer. Quality checking and MAG completeness were assessed using CheckM(*84*). Genomic annotations were performed using rapid prokaryotic genome annotation (Prokka v1.14.6), the database of Clusters of Orthologous Groups (COG) and the Kyoto Encyclopedia of Genes and Genomes (KEGG). Pangenomic analysis were performed using Anvi’o version 7.1(*85*), to compare MAG-CC1 with other 10 high-quality *Endozoicomonas* genomes retrieved from public databases (Supplementary Table S6).

### Bioinformatic analysis for amplicon sequencing

QIIME2 (v.2022.2)(*86*) was used to import the sequencing reads of the 16S rRNA gene amplicons, with primers trimmed using the q2-cutadapt command. Forward and reverse reads were truncated at positions 132 and 130, respectively. Amplicon sequence variants (ASVs) were denoised and clustered using DADA2(*87*) implemented in QIIME2. Taxonomy was inferred using q2-feature-classifier classify-sklearn and the SILVA database v138.1, with phylogenetic trees constructed using q2-align-to-tree-mafft-fasttree. Phyloseq objects were exported from QIIME2 data, including the feature table, taxonomy table, rooted tree, and metadata, for further analysis in R (version 4.1.2). An alpha-diversity index (Pielou’s evenness) was generated using rarefied data. Normality was tested using the Shapiro-Wilk test, and the Analysis of Variance (ANOVA) was used to assess statistical differences in the index (PAST 4.03). Raw ASV data were normalized using DESeq2(*88*) and used to generate taxa barplots and analyze species indicators and networks. The same normalized data were used to calculate Sørensen dissimilarity, which was visualized via Principal coordinates analysis (PCoA), using the phyloseq package implemented in R. Permutational multivariate analysis of variance (PERMANOVA) was performed on the data matrix(*89*) to compare the composition of the microbial communities. Permutational multivariate analysis of dispersion (PERMDISP) was also performed, with 999 permutations, using the betadisper function (implemented in vegan 2.6-4).

### Species indicator

IndicSpecies (v. 1.7.14) implemented in R was used to identify those ASVs that are indicators for sample groupings. We used func = “IndVal.g” and control with number of permutations = 9999. We filtered the data by excluding ASVs with p-values ≥ 0.05 to improve accuracy and reduce the excess of ASVs. Matrices were exported for plotting and analyzing in Cytoscape. The indicator value index (IndVal or Stat) is the product of two components, “specificity” and “fidelity”. These values range from 0.00 to 1.00. A specificity value = 1.00 means that a given ASV was found in all samples from a sample group. A fidelity value = 1.00 means that a given AVS was found exclusively in that sample group. Thus, a Stat value ≅ 1.00 indicates high specificity and high fidelity.

### Molecular and morphological taxonomy of hosts

Sequences of the COI gene from each host (corals and crinoid) were trimmed manually. Alignment of COI gene sequences and the contigs were assembled in BioEdit 7.2. After performing preliminary analysis, BLASTN analysis was conducted against the NT database at the National Institute of Biotechnology Information (NCBI) to identify the hosts.

Due to the low identity of the crinoid sequences against the NT database, the specimens were also air-dried to observe morphological features, using a stereomicroscope (ZEISS SteREO Discovery V20). The specimens were identified based on diagnostic characters in keys and species descriptions in the historical literature and taxonomic reviews for the family Thalassometridae and subsequently for the genus *Koehlermetra* (for example, Carpenter(*29*), Clark(*30*)(*29–33*)). Crinoid classification and images of the type material available in the WoRMS database(*90*) were also used for comparison with the material. Following these results, the crinoid COI gene sequences were compared to the *K. porrecta* COI gene, retrieved from ENA under accession number KC626561.1.

### Phylogenetic analysis

The Bacterial Genome Tree Service(*91*) was used to compute the phylogenomics for *Endozoicomonas* sp. MAG-CC1 and 10 other high-quality *Endozoicomonas* spp. genomes from public repositories (Supplementary Table S6), using the codon tree method, which consists of aligning 483 single-copy genes and their translated proteins to infer phylogenetic trees. The tree was rooted using *Halomonas meridiana* NBRC 15608 as an outgroup. In addition, we employed the Genome-to-Genome Distance Calculator 3.0(*91*, *92*) to perform a comprehensive analysis of the digital DNA-DNA hybridization (dDDH) and assess the degree of similarity between the MAG-CC1 and its closest reference genome. The average nucleotide identity (ANI) based on BLAST+ was measured using JSpecies Web Server(*93*). In addition, the average amino acid identity (AAI) was calculated using EzAAI(*94*).

### Fluorescence in-situ hybridization and scanning electron microscopy

To visualize the cellular localization of the targeted microorganisms in coral tissue and crinoid stalk and crown cells, we performed fluorescence *in situ* hybridization (FISH) in sections of selected samples. Similarly, to visualize superficial structures, polymeric extracellular structures and microbial cells colonizing corals and crinoids, we performed scanning electron microscopy (SEM).

For FISH, all samples were fixed immediately after collection in 4% paraformaldehyde overnight at 4 °C, and then rinsed in a solution of PBS:EtOH (1:1, v:v) for 5 min. Samples were stored in PBS at 4 °C. In the laboratory, coral samples were embedded in 1.5% agarose and decalcified by subsequent baths in 20% EDTA (samples kept at 4 °C) twice a week until skeletons were no longer visible. Tissue samples were then washed three times in PBS. All samples were dehydrated in an EtOH series (30, 50, 70, 90, 3x 100%) and embedded in LR White resin at 4 °C (1:1 EtOH:LR White overnight, 2x 100% LR White for 1 h each). Samples were polymerized in BEEM capsules at –20 °C under UV light for 5 days. Thin sections were made and collected on microscope slides. For hybridization, sections were first treated with a solution composed of 50 µg/ml proteinase K + 20 mM Tris HCl buffer, to permeabilize cell membranes. Sections were then covered with a hybridization buffer (formamide 30% + 0.9 M NaCl + 20 mM Tris HCl + 0.01% SDS) and probes were added, to a final concentration of 25 ng/µl. The tissues were hybridized using probes specific for *Endozoicomonas* spp.(*35*, *94*) (Endozoi663, 5’-Alexa594 GGA AAT TCC ACA CTC CTC-3’ and Endozoi736, 5’-Alexa594 GTC AGT GTC AGA CCA GAG-3’) and Crenarchaeota/Thaumarchaeota (AOA)(*36*) (Cren512, 5’Alexa647-CGG CGG CTG ACA CCA G-3’) as well as a non-sense NonEUB(*95*) (5’-Alexa532 ACT CCT ACG GGA GGC AGC-3’) and NonARCH(*96*) (5’-Alexa532 CAC GAG GGG GCG GTT AAG GA-3’) probes (Supplementary Table S7). Additional FISH experiments (attached to Supplementary Data) were performed using the fluorophore Alexa637 for probing Endozoi663, Endozoi736 and Cren512, in addition to Alexa637 for probing the Antisense NonEUB and NonARCH. While specific probes for *Nitrosopumilaceae* were not available in the literature, our decision for using a probe designed for most Crenarchaeota/Thaumarchaeota was justified by the factthat ASVs assigned to the *Nitrosopumilaceae* constitute 89.52% of all archaeal ASV in our data. Samples were protected from light and incubated at 46 °C for 1.5 h. Sections were then washed in a washing buffer (0.112 M NaCl + 20 mM Tris HCl + 0.01% SDS + 5 mM EDTA) at 48 °C for 10 min and rinsed in cold RNAse/DNAse-free water to remove remaining salts. Finally, sample slides were dried and mounted in Fluoroshield + Dapi (Sigma-Aldrich). All samples were observed in a Leica SPE Confocal Laser Scanning microscope. And fluorescence stack images were processed using ImageJ in order to create Z-maximum images and 3D videos. Ten sections (n = 10) for each host replicate were observed.

For SEM analysis, samples were fixed immediately after collection, in a solution composed of 2.5% glutaraldehyde + 4% paraformaldehyde + 0.1 M sodium cacodylate buffer overnight at 4 °C and then rinsed 3x in 0.1 M sodium cacodylate buffer. In the laboratory, samples were dehydrated in an EtOH series (30, 50, 70, 90, 3x 100%), critical-point dried in a Baltec CPD030 dryer, mounted on stubs, and coated with gold. Samples were observed using a Zeiss EVO 10 Scanning Electron Microscope.

## Acknowledgments

The authors would like to thank Instituto de Pesquisas Jardim Botânico do Rio de Janeiro (JBRJ) for the Confocal Laser Scanning microscope use.

## Author contributions

Conceptualization: FM, RP, TT.

Methodology: FM, RP, TT, AG, LH, GD, AB, RBM.

Analysis: FM, RP, TT, AG, LH.

Visualization: FM, LH.

Supervision: RP, TT.

Writing—original draft: FM, RP, TT, AG, LH, GD, AB, RBM, VP.

Writing—review & editing: FM, RP, TT, AG, LH, GD, AB, RBM, VP.

## Data and materials availability

Raw amplicon-sequencing and MAG-CC1 data are available on NCBI’s Read Archive under accession PRJNA991147. *Endozoicomonas promiscua* sp. nov. was registered on SeqCode and the nomenclature details can be checked under the Canonical URL <https://seqco.de/i:49122>. The complete bioinformatic pipeline including scripts for figure reproduction is available through the GitHub repository at <https://github.com/modolon/deep-sea_crinoid_corals>.

## Supplementary Materials

### Supplementary Figures

**Supplementary Figure S1.** Sørensen dissimilarity for *D. pertusum* and *S. variabilis* associated with crinoids and those not associated with them. *D. pertusum*, p-value = 0.400; for *S. solenosmilia*, p-value = 0.812; PERMANOVA.

**Supplementary Figure S2.** Correlation analysis (Spearman correlation) between 15 most abundant ASV families and nitrogen forms (nitrate, nitrite and ammonium) and dissolved organic carbon (DOC). Prv. and Abd. indicates the prevalence and abundance of the taxa. Dendograms clusterized the rows (taxa) and columns (factors) by similarity

**Supplementary Figure S3**. Individualized fluorescent channels used in FISH observation using Endozoicomonas spp. (left panel) and AOA probes (right panel).

**Supplementary Figure S4**. Fluorescent channels used in FISH observation using probes for Endozoicomonas spp. (left panel) and AOA (right panel).

### Supplementary Tables

**Supplementary Table S1.** Taxonomic classification of deep sea corals and crinoids by CO1 sequencing.

**Supplementary Table S2.** Summary reads statistics from amplicon sequencing and ASV table with raw read count.

**Supplementary Table S3.** Genomic features from annotation of MAG-CC1 using Prokka pipeline.

**Supplementary Table S4.** Set of genes retrieved from pangenomic analysis of *Endozoicomonas* genus.

**Supplementary Table S5.** Metadata related to sampling sites.

**Supplementary Table S6.** Genome set and accession IDs used for phylogenomic and pangenomic analysis.

**Supplementary Table S7.** List of probes used for fluorescence in situ hybridisation (FISH).

