## Supplementary Information for "Promiscuous endosymbionts in deep-sea corals and crinoids are shaped by nitrogen cycling"

### Content

**Supplementary Figure S1.** Sørensen dissimilarity for *D. pertusum* and *S. variabilis* associated with crinoids and those not associated with them. *D. pertusum*, p-value = 0.400; for *S. solenosmilia*, p-value = 0.812; PERMANOVA.
**Supplementary Figure S2.** Correlation analysis (Spearman correlation) between 15 most abundant ASV families and nitrogen forms (nitrate, nitrite and ammonium) and dissolved organic carbon (DOC). Prv. and Abd. indicates the prevalence and abundance of the taxa. Dendograms clusterized the rows (taxa) and columns (factors) by similarity
**Supplementary Figure S3**. Individualized fluorescent channels used in FISH observation using Endozoicomonas spp. (left panel) and AOA probes (right panel).
**Supplementary Figure S4**. Fluorescent channels used in FISH observation using probes for Endozoicomonas spp. (left panel) and AOA (right panel).

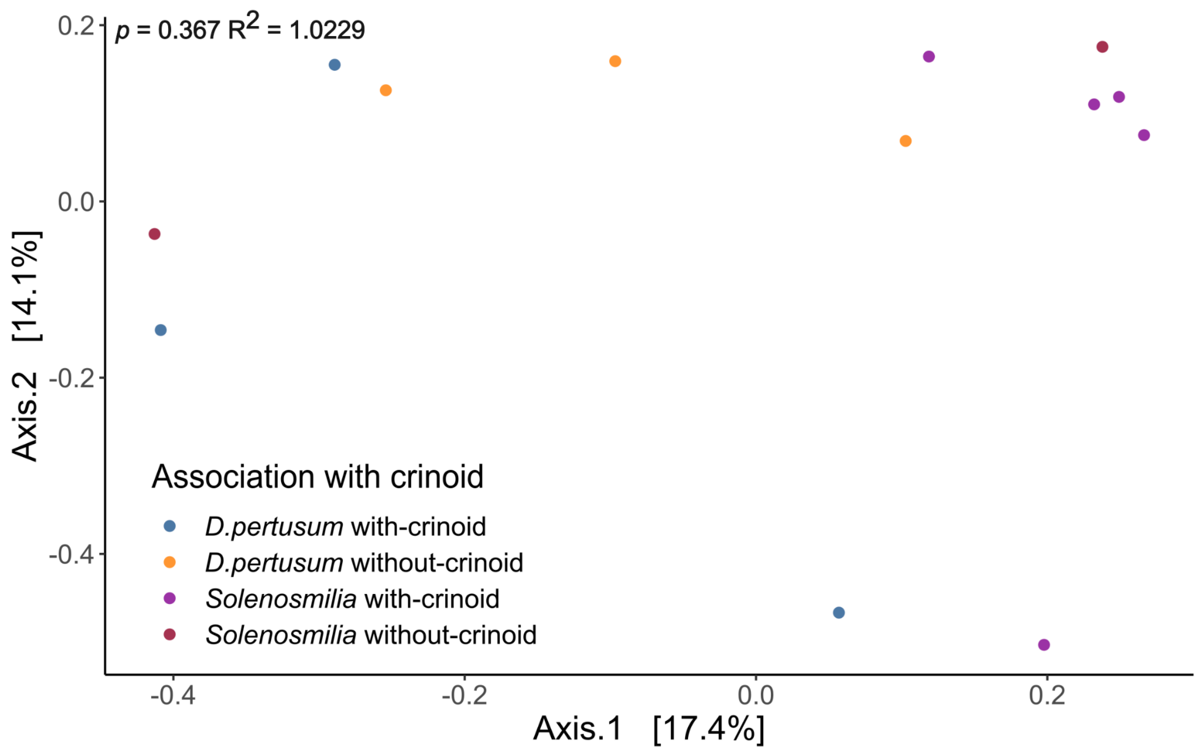


**Supplementary Figure S1.** Sørensen dissimilarity for *D. pertusum* and *S. variabilis* associated with crinoids and those not associated with them. *D. pertusum*, p-value = 0.400; for *S. solenosmilia*, p-value = 0.812; PERMANOVA.

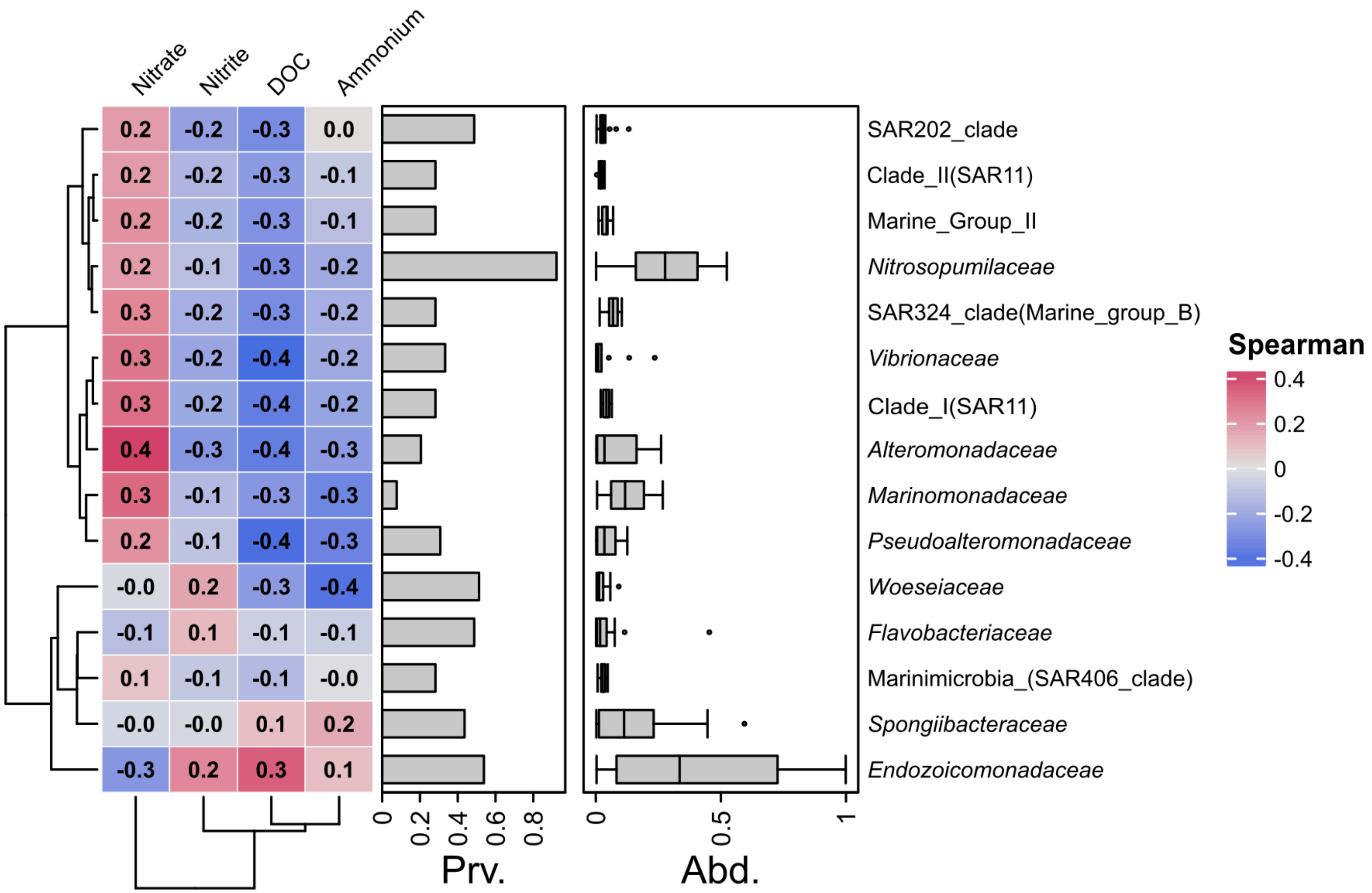


**Supplementary Figure S2.** Correlation analysis (Spearman correlation) between 15 most abundant ASV families and nitrogen forms (nitrate, nitrite and ammonium) and dissolved organic carbon (DOC). Prv. and Abd. indicates the prevalence and abundance of the taxa. Dendograms clusterized the rows (taxa) and columns (factors) by similarity

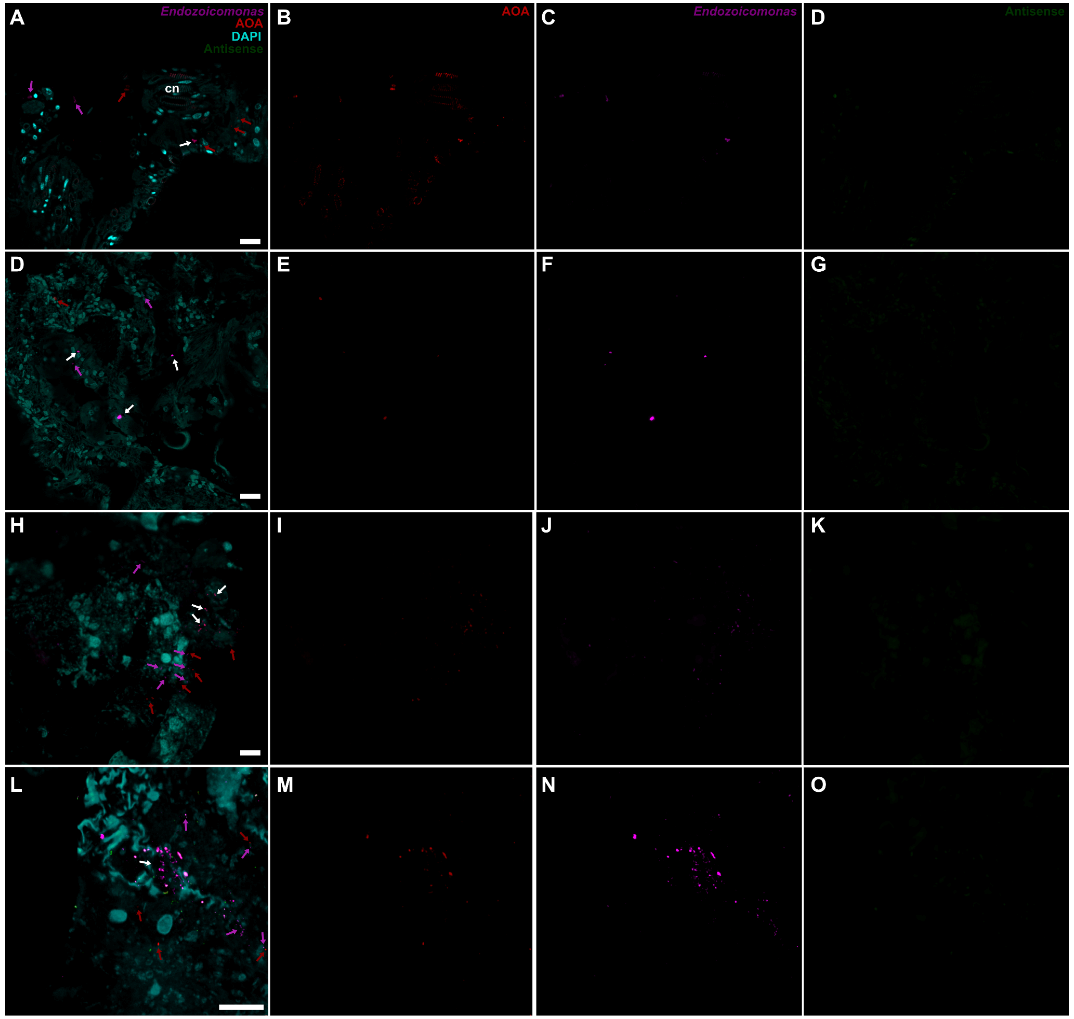


**Supplementary Figure S3**. Individualized fluorescent channels used in FISH observation using Endozoicomonas spp. and AOA probes. Fluorophore Alexa637 was used for *Endozoicomonas* probes (Endozoi663 and Endozoi736). Alexa637 also applied for AOA. Alexa594 for Antinse probes (NonEubac and NonArch).

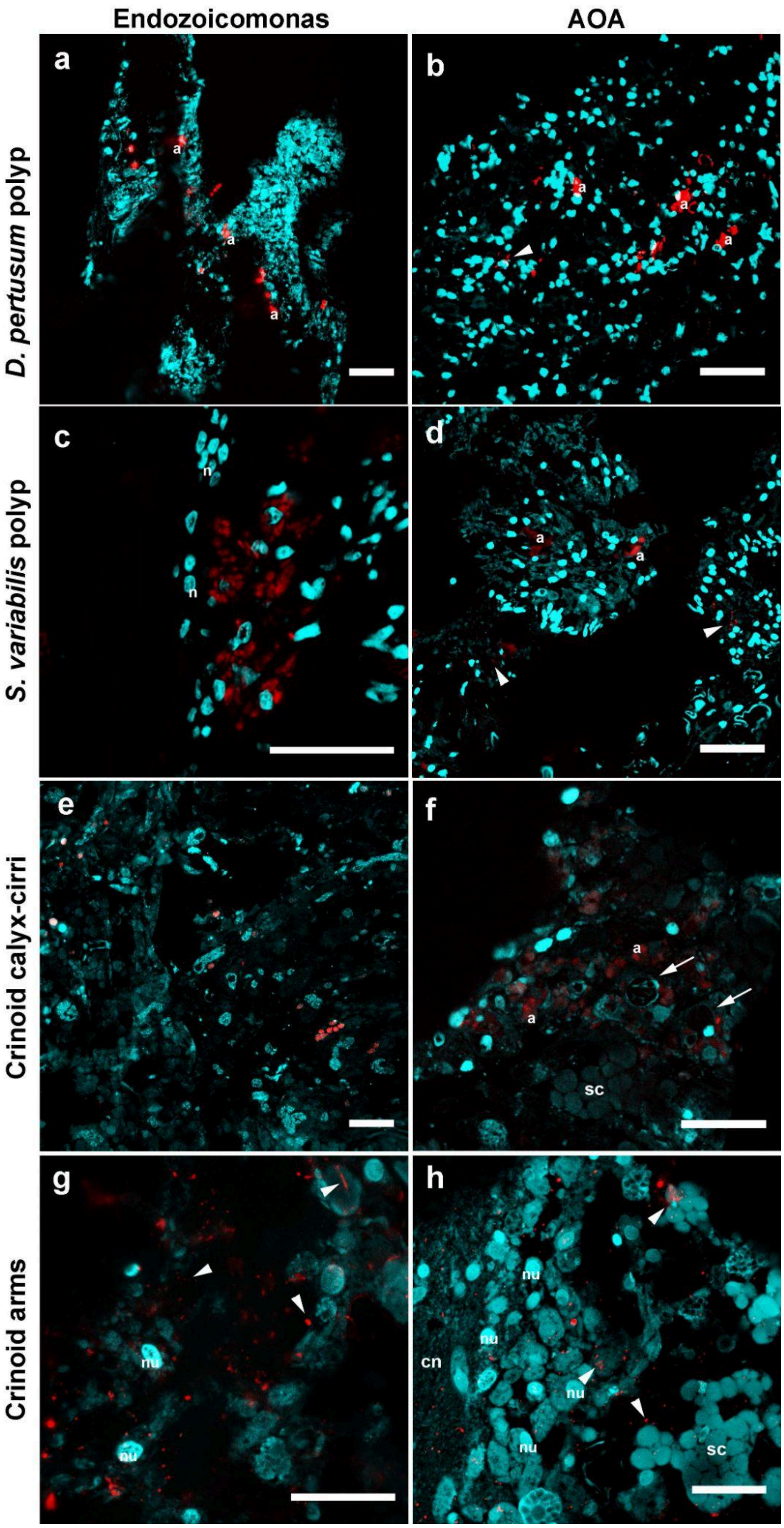


**Supplementary Figure S4**. Fluorescent channels used in FISH observation using probes for Endozoicomonas spp. (left panel) and AOA (right panel). Fluorophores Alexa637 was used for all probes. Bars = 5 μm.

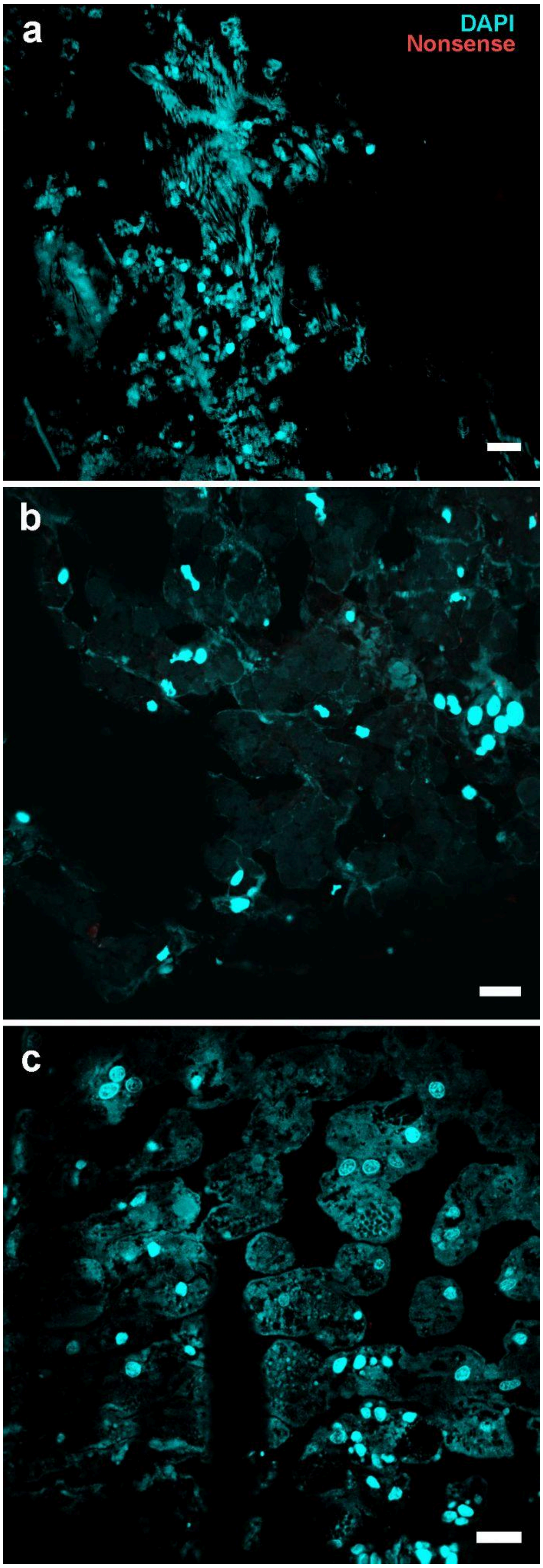


**Supplementary Figure S5**. FISH images using NonSense (antisense) probing (Non-Eubac) to test for probe specificity in samples from *Desmophyllum pertusum* tissue (A), *Solenosmilia variabilis* tissue (B) and Crinoid calyx-cirri tissue (C). In cyan-blue is DAPI and in red the non-sense probe (fluorophore Alexa637). Bars = 10 µm.
